# The Hill-type equation reveals the regulatory principle of target protein expression led by p53 pulsing

**DOI:** 10.1101/2023.10.24.563713

**Authors:** Xiaomin Shi

## Abstract

The central dogma indicates the basic direction of gene expression pathways. For activated gene expression, the quantitative relationship between various links from the binding of transcription factors (TFs) to DNA to protein synthesis remains unclear and debated. There is consensus that at a steady state, protein levels are largely determined by the mRNA level. How can we find this steady state? Taking p53 as an example, based on the previously discovered Hill-type equation that characterizes mRNA expression under p53 pulsing, I proved that the same equation can be used to describe the average steady state of target protein expression. Therefore, at steady state, the average fold changes in mRNA and protein expression under TFs pulsing were the same. This consensus has been successfully demonstrated. For the p53 target gene *BAX*, the observed fold changes in mRNA and protein expression were 1.40 and 1.28, respectively; the fold changes in mRNA and protein expression calculated using the Hill-type equation were both 1.35. Therefore, using this equation, we can not only fine-tune gene expression, but also predict the proteome from the transcriptome. Furthermore, by introducing two quantitative indicators, we can determine the degree of accumulation and stability of protein expression.

## Introduction

According to the central dogma of molecular biology, the gene expression pathway is from DNA to RNA to protein (1, 2), which is the process of transcribing gene-encoded information into mRNA and translating it into functional proteins. For gene expression, mRNAs and proteins provide useful readouts that connect genes and phenotypes (3). Generally, there are two different gene expression pathways involved. One pathway is the basal pathway, which regulates homeostatic expression, and the other pathway is the activated or repressed pathway, which dynamically regulates expression in response to environmental stimuli (4-8).

For basal gene expression, steady-state protein levels depend on the transcription rate, mRNA half-life, translation rate constant, and protein half-life (9, 10). Moreover, situations with high transcription rates and low translation rate constants should be ruled out (10). Under environmental stimuli, for gene expression activated by transcription factors (TFs), studies involving transcripteomics and proteomics indicate that the protein level at steady state is largely contributed by the mRNA level (3, 11, 12), however, there is little relationship between the protein level and synthesis or protein half-life.

Here, the steady state can be defined as a relatively long-term process experienced by cells (11). Therefore, both cell proliferation and differentiation can be considered steady states (11). Apoptosis and senescence also occur in a steady state (13, 14). The steady state achieved after stimulation differs from that before perturbation (15).

For the steady state before stimulation in dendritic cells, mRNA levels explain 68% of the change in protein expression, and for the approximate steady state after stimulation with lipopolysaccharide, mRNA levels explain 90% of the change in protein expression (12, 16). In addition, during the development of *C elegans*, the fold changes in mRNA and protein expression were almost identical. The transcript fold change was 2.02, and the protein level was 2.05 (17). Therefore, we can speculate that the fold-changes in mRNA and protein expression in the steady state are equal. Could we theoretically determine this special steady state of protein expression? Which factors determine protein expression levels in the steady state?

Gene expression under environmental stimuli is driven by TFs, which map the corresponding stimulus. TF dynamics encode specific stimuli (18). The binding components of TF and DNA constitute the decoder of TF dynamics. This initiates the expression of the corresponding target gene and fulfils the relevant function. Some TFs exhibit oscillatory dynamic behaviors, in which the duration, frequency, and amplitude can all encode components of gene expression, thereby leading to different cell fates (19, 20).

The tumor suppressor p53 is the most extensively studied TF. In response to DNA damage, p53 concentrations exhibit a series of pulses of fixed duration, frequency, and amplitude, whereas the number of p53 pulses increases with the degree of DNA damage caused by *γ* irradiation (21). Changing p53 dynamics from pulsed to sustained behavior leads to an altered cell fate from DNA repair to senescence (22). For gene expression driven by p53 pulsing, there is an interesting phenomenon in which the levels of mRNA and target protein expression are very similar. For example, for the *MDM2* gene, 10 h or 24 h after stimulation, the fold change in mRNA was 2.2 (10 h)(23) or 2.0 (24 h)(24), while the protein expression was 1.8 (24 h), which can be regarded as the average steady state over the cell population (23). General speaking, TFs dynamics is pulsed or sustained. In fact, the analytical results always face to all TFs dynamics. p53 dynamics has been studied for more than 25 years, and there is some data we may use to verify the analytical results. Therefore, without loss of generality, I will use target gene expression under p53 pulsing as an example to determine the principle governing target protein expression at a steady state.

The classic Hill equation originated from physiology, and became a basic equation in biochemistry. Physiology is the precise quantitative study of various dynamical processes within an organism from a macro perspective, therefore, the present findings may help integrate TFs dynamics into systems physiology.

Following the genetic information flow specified by the central dogma, we obtained a modified Hill equation to characterize the average p53 DNA-binding probability (25). I also found a Hill-type equation that could predict the fold-change in target mRNA expression under p53 pulsing (26).

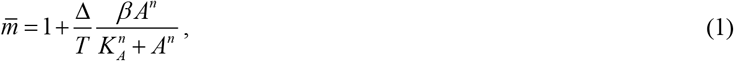

where Δ denotes the duration, *T* is the period, *A* is the amplitude, *β* is the maximal fold change in mRNA expression, and *K*_*A*_ is the dissociation constant.

Here, I will complete the last step in describing the central dogma, from mRNA to protein. In response to pulsed p53 input dynamics, p21 mRNA dynamics show pulsing expression; however, p21 protein dynamics exhibit rising expression (27). The half–life of mRNA and protein determines the stability of expression dynamics (23, 24, 28). However, the relationship between steady-state mRNA and protein expression levels remains unclear. Therefore, I tried to prove that the average steady-state fold changes in mRNA and protein expression under p53 pulsing were equal, *i*.*e*., the average protein fold change at steady state was equal to *m*.

## Methods

### Mathematical model of p53 target protein expression dynamics and its analytical solution

To achieve this goal, we must develop a useful and accurate mathematical model for gene expression under pulsed TF dynamics (16, 29). Some models do not include the basal transcription term,(30) but some early models provided necessary terms for environmental stimuli to activate gene expression.(31) The basal transcription term is necessary for introducing fold change. The ordinary differential equation for mRNA dynamics is (26)

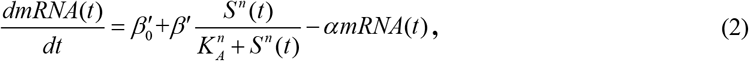

where

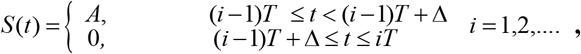

*mRNA*(*t*) denotes the target mRNA concentration, *S* (*t*) is the square wave function of the TF p53 pulses, 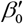 is the basal transcription rate, *β* ′ is the maximal transcription rate activated by p53, *α* is the mRNA decay rate constant, and *n* is the Hill coefficient. *T* is the period, *A* is the amplitude, which are the same as described in Equation 1. The initial condition is *mRNA*(0) = *m*_0_, where *m*_0_ denotes the basal mRNA concentration.

The equation for the protein expression dynamics is (9, 10, 23)

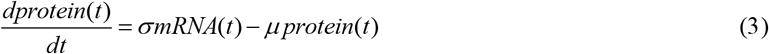

where *protein*(*t*) represents the target protein concentration, and *σ* and *μ?*denote the rate constants of translation and degradation, respectively. The initial protein concentration was determined using *protein*(0) = *p*_0_.

For activated expression dynamics, considering 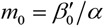, *p*_0_ = *σ m*_0_ *μ*, letting become *m*(*t*) = *mRNA*(*t*) *m*_0_, *P*(*t*) = *protein*(*t*) / *p*_0_, Equation 2 and Equation 3 become

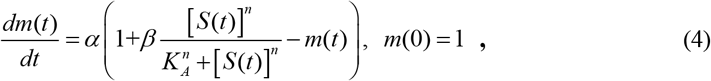

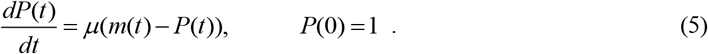

where *m*(*t*) and *P*(*t*) represent the fold changes in mRNA and protein expression, respectively, and 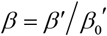 denotes the maximal fold change in transcription. It seems that I have never seen gene expression levels appear in a dimensionless form in the differential equation. This is a crucial step. According to equation 5, the steady-state levels of mRNA and target proteins are the same. Because Equation 4 has been solved, the solution from Shi (26) is rewritten as

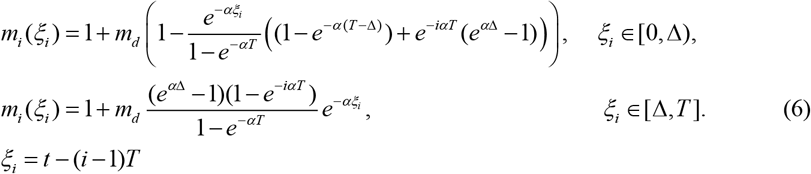

where

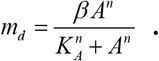

*m*_*i*_ (*ξ*_*i*_) represents the mRNA fold change during the i-th TF pulse.

Solving Equation (5) is challenging. Salazar et al. (32) provides a method for solving such equations. Assuming *α* ≠ *μ*, we can obtain analytical solutions for target protein expression in response to p53 pulsing as follows:

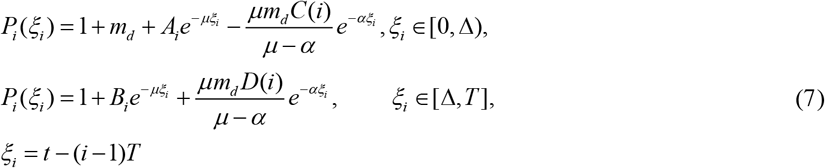

*P*_*i*_ (*ξ*_*i*_) represents the protein fold-change under the i-th p53 pulse. Here, a detailed derivation of *A*_*i*_ and *B*_*i*_ is presented in Appendix. The detailed solving process for Equation 5 is provided in Appendix.

## Results

### Steady-state fold changes in target protein expression exhibits oscillations

The steady state is the relatively long-term behavior of the cells. Neither Salazar el al. (32) nor Shi et al.(25) conducted this steady-state analysis. Shi (26) began this type of analysis. By inspecting Equation 7, for sufficient p53 input pulses, letting *i* →∞, we can obtain the steady state of the target protein expression dynamics:

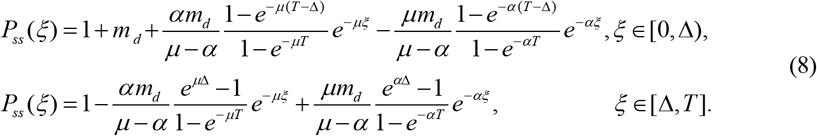

*P*_*ss*_ (*ξ*) reached its maximum and minimum at Δ and 0 or *T*, respectively. Therefore, the steady-state of target protein expression dynamics is a repetitive and invariant oscillation. The maximal fold change or peak of oscillations was

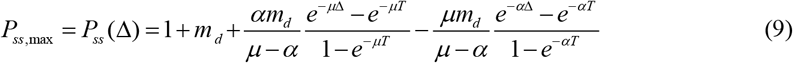

The minimal fold change or valley is

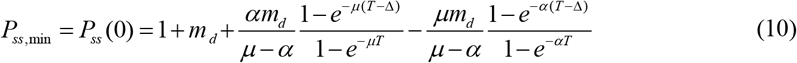

For a given target protein, *β* and *K*_*A*_ are relatively fixed, and the amplitude depends on the p53 dynamic parameters and the half-life of the mRNA and protein. Apparently, the smaller the amplitude is, the more stable the steady state.

### A Hill-type equation can characterize the constant steady state

Let us now examine the characteristics of the steady state within the limits of *α* and *μ*. From Equations A6 and A7, when *αT* ≪ 1, *μT* ≪ 1was applied,

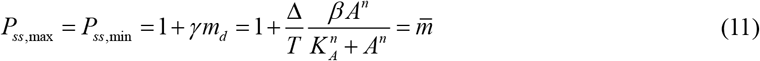

Equation 11 is the same as Equation 1, which is exactly the Hill-type equation found previously (26).

Similarly, when *αT* ≪ 1, *μ*Δ ≫ 1, *i*.*e. μT* ≫ 1, there are

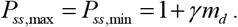

When *μT* ≪ 1, *α*Δ ≫ 1, *i*.*e. αT* ≫1, there are

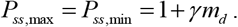

When *α*Δ ≫ 1, *μ*Δ ≫ 1, *i*.*e. αT* ≫ 1, *μT* ≫ 1, there are

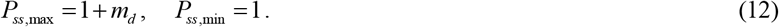

Equation 12 is the same as the classical Hill equation, which governs the steady-state mRN A fold-change under sustained p53 dynamics (26). Therefore, in the limit of a very long or short half-life for mRNA and protein, oscillations contract into a constant line that is very stable.

For several target genes of p53, Table 1 lists the results for *α, μ, β*, and *K*_*A*_. Only 3 genes had complete data. *α* and *μ* determine the trajectory of target protein expression dynamics (23). The four scenarios discussed above correspond to the four sets of protein expression dynamics defined in Hanson et al. (23).

**Table 1.** The observed values of *α, μ, β* and *K*_*A*_ corresponding to genes

| Gene | Function | mRNA decay rate constant | Protein degradation rate constant | Dissociation constant | Maximal mRNA fold change |
| --- | --- | --- | --- | --- | --- |
| *PUMA* | Apoptosis | 0.73 h<sup>-1</sup> | 0.056 h<sup>-1</sup> | 260 nM | Unknown |
| *MDM2* | Feedback inhibition | 0.27 h<sup>-1</sup> | 0.7916 h<sup>-1</sup> | 12.3 nM | $2^{2.7861} = 6.898$ |
| *p21* | Cell cycle arrest | 0.265 h<sup>-1</sup> | 0.2546 h<sup>-1</sup> | 4.9 nM | $2^{3.9008} = 14.937$ |
| *BAX* | Apoptosis | 0.018 h<sup>-1</sup> | 0.0262 h<sup>-1</sup> | 73 nM | $2^{1.2373} = 2.358$ |
| *GADD45A* | DNA repair | Unknown | Unknown | 7.7 nM | $2^{3.1949} = 9.157$ |

As shown in Table 1, the mRNA decay rate constants of *PUMA, MDM2*, and *p21* are from (28), and the mRNA half-life of *BAX* is 38.77 h (33), thus, the mRNA decay rate is *α* = ln 2 38.77 ≈ 0.018 h^−1^ (23). The protein degradation rate constants of *PUMA, MDM2, p21*, and *BAX* were obtained from (23). The dissociation constants of *PUMA, MDM2, p21, BAX*, and *GADD45A* are from (34). For maximal mRNA expressions of *p21*(*CDKN1A*), *GADD45A, MDM2*, and *BAX*, log_2_ (maximal mRNA fold change) were 3.9008(3 h), 3.1949(3 h), 2.7861(3 h), and 1.2373(9 h)(24), thus, the maximal fold changes were 14.937, 9.157, 6.898, and 2.358, respectively.

### The average fold changes in mRNA and protein expression were the same at steady state

The average fold change of the target protein over the i-th period can be calculated as follows (Appendix) :

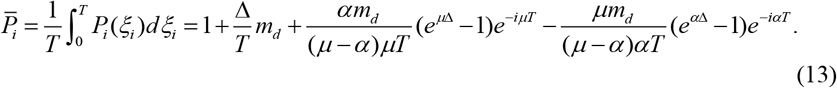

Letting *i* →∞, the average steady state is

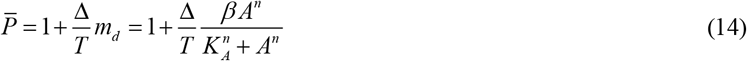

As we expect

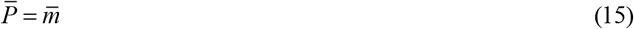

Therefore, we proved that at steady state, the average fold changes in mRNA and protein expression were the same. This result also helds for the average levels of different proteins and their corresponding mRNAs within a single cell, which is consistent with the observed results (17). Similarly, for the average of the same proteins across the cell population, the average mRNA and protein expression levels were also the same.

Next, we considered p53 target gene *BAX* as an example to examine the predictability of the Hill-type equation for protein expression. The observed fold-change in the expression of the BAX protein 24 h after stimulation was 1.28 (23), and that of the BAX mRNA were 1.40 (28) or 2^0.7802^ = 1.72 (24). For any cell, assuming that Δ *T* remains unchanged, the average BAX expression over the cell population can be calculated using the Hill-type equation (26):

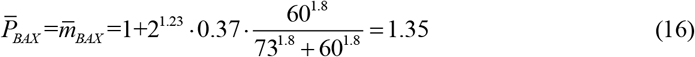

The values for *β* and *K*_*A*_ are listed in Table 1. The Hill coefficient is *n* = 1.8 (34). *A* = 60 nM (25). The duty cycle of p53 pulsing *γ* can be written as

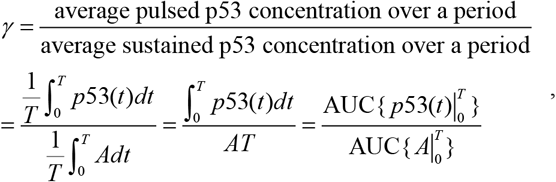

where 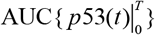represents the area under the curve *p*53(*t*). As shown in Fig. 1, for a general periodic function, *p*53(*t*), during [0,*T*] can be approximated by a square wave function *S* (*t*), as long as

**Figure 1.**
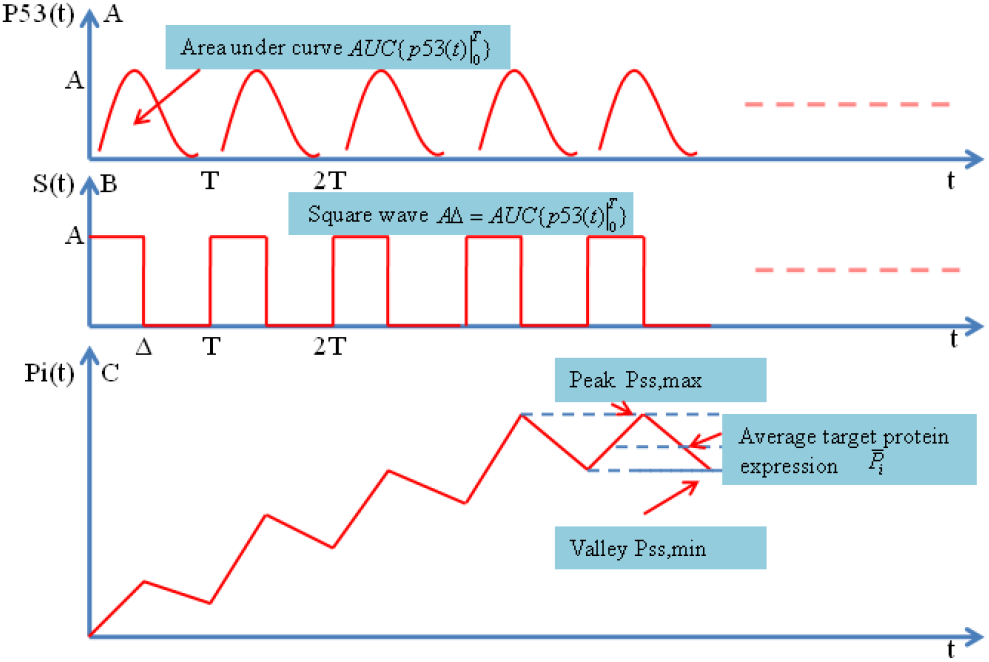
Target protein expression led by p53 pulsing. A. p53 pulses with amplitude A and period T. For each pulse, the area under curve is denoted by 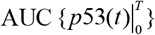. B. Each p53 pulse is approximated by square wave with duration 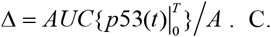 After several pulses, a typical rising target protein expression dynamics attains the steady state which has the same peaks and valleys. The peak-to-valley ratio can describe the stability of target protein expression dynamics.

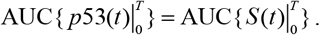

Thus,

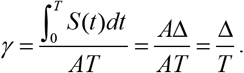

Here, *γ* is taken as 0.37 to minimize the error between the predicted p21 mRNA fold change and the observed fold change(26).

### A longer mRNA or protein half-life determines the relaxation time of target protein expression dynamics

We previously obtained the relaxation time for mRNA dynamics (26). Similarly, the time of protein dynamic trajectory relaxation to the average steady state can be calculated by (25, 26, 32)

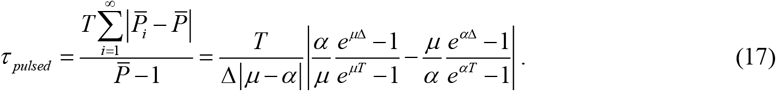

When *αT* ≪1, *μT* ≪ 1, *τ* _*pulsed*_ can be expanded in the Taylor series:

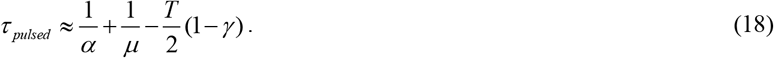

Thus, the number of p53 pulses required to reach the average steady state is

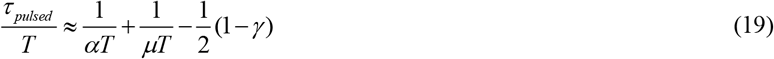

For sustained p53 input dynamics, Δ = *T*, thereby

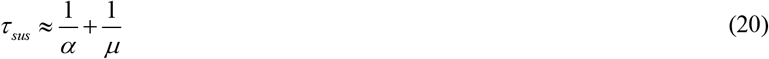

Therefore, compared to sustained input, pulsed p53 dynamics cause target protein dynamics to reach maximum more quickly. In other words, if the half-lives of mRNA and protein and *T* (1−*γ*) 2 have the same order of magnitude, oscillatory p53 input enhances the sensitivity of protein expression. On the other hand, as shown in Table 1, the gene encoding the *BAX*, which has a longer half-life of mRNA and protein requires multiple p53 pulses to reach a steady state, which not only provides sufficient time for DNA repair but also leads to the accumulation of sufficient expression levels required for triggering apoptosis.

### The index of target protein accumulation under multiple p53 pulses

To understand the degree of accumulation of the target protein in response to multiple p53 input pulses, the index of protein accumulation can be defined as

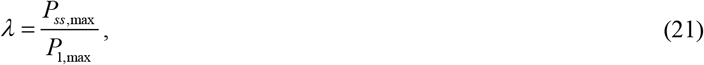

which is the ratio of the maximal protein fold change at steady state to the maximal protein fold change during the 1st pulse.

*P*_*ss*,max_ is given by Equation 9, and

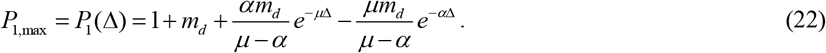

When *αT* ≪1, *μT* ≪ 1, there are

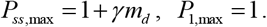

Thus

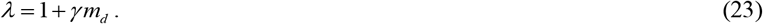

Similarly, when *αT* ≪1, *μ*Δ ≫ 1, *i*.*e. μT* ≫ 1, there is

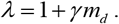

When *α*Δ ≫ 1, *i*.*e. αT* ≫ 1, and *μT* ≪ 1, there is

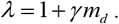

When *α*Δ ≫ 1, *i*.*e*., *αT* ≫1and *μ*Δ ≫ 1, *i*.*e*.*μT* ≫ 1, there are

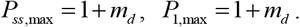

Thus

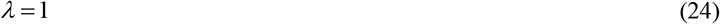

Therefore, proteins with longer half-lives have a higher accumulation; however, proteins with shorter half-lives have a lower accumulation.

The first three cases discussed above broadly correspond to the rising expression dynamics observed in Hanson et al. (23), and the last corresponds to the oscillatory expression dynamics. In other words, the rising expressions have higher accumulation, and the pulsing expressions have lower accumulation.

### The peak-to-valley ratio can measure the stability of mRNA and protein expression dynamics under multiple p53 pulses

According to Equation 6, the mRNA expression dynamics at steady state also exhibit oscillatory behavior. Let, The steady state for mRNA dynamics can be written as

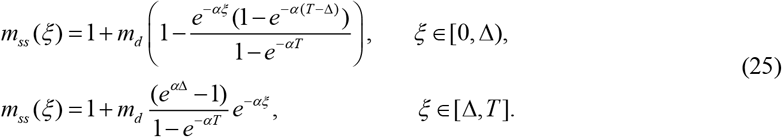

The peak is

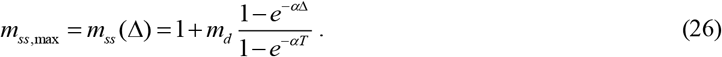

And, the valley is

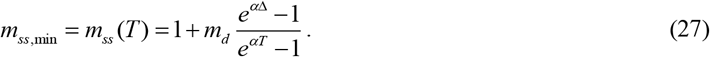

For a given gene, the peak-to-valley ratio for target mRNA expression can be defined as

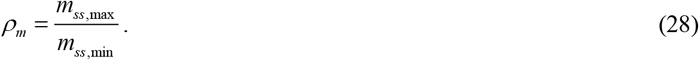

The closer *ρ*_*m*_ is to 1, the more stable the mRNA expression dynamics are. In particular, when *αT* ≪ 1

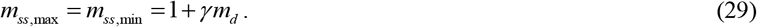

Thus

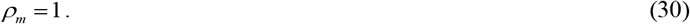

Therefore, the longer the mRNA half-life is, the more stable the mRNA expression dynamics are. Similarly, when *α*Δ ≫ 1, *i*.*e. αT* ≫1,

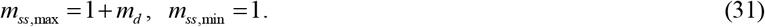

Therefore,

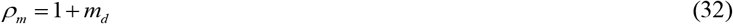

Therefore, target mRNA dynamics with shorter half-lives are unstable.

Similarly, we can define the peak-to-valley ratio that characterizes the stability of the target protein expression dynamics as:

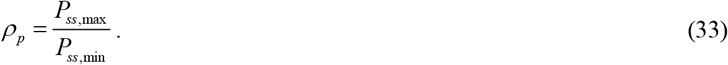

As seen from the previous discussion, for the three extreme cases of *α* and *μ, ρ*_*p*_ = 1. However, for the short half-lives of mRNA and target protein, according to Equation 12, *ρ*_*p*_ = 1+ *m*_*d*_ > 1, therefore, the target protein is unstable in this case. When mRNA has a short half-life, *ρ*_*m*_ = 1+ *m*_*d*_, therefore, translation of mRNA does not improve its stability. By calculating the peak-to-valley ratio of each target gene, we can compare the stability of the expression of different genes.

The index of mRNA accumulation can be defined as

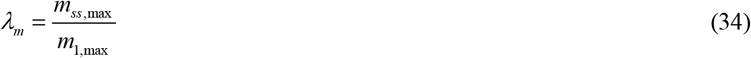

From Equation 6, we obtain

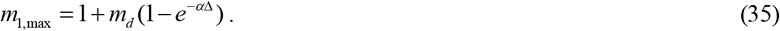

For *αT* ≪1, *λ*_*m*_ = 1+ *γ m*_*d*_, and for *αT* ≫1, *λ*_*m*_ = 1. Therefore, stable mRNA has a high degree of accumulation.

As shown in Table 2, for the four combinations of half-lives, both long mRNA and short protein half-lives, as well as short mRNA and long protein half-lives can produce stable proteins. Only short mRNA and short protein half-lives produce unstable proteins. Stable proteins always have high accumulation.

**Table 2.** The four combinations of mRNA and protein half-lives determine protein stability

| mRNA half life | Protein Half life | mRNA stability, $\rho_m$ | Protein stability, $\rho_n$ | mRNA accumulation, $\lambda_m$ | Protein Accumulation, $\lambda_n$ |
| --- | --- | --- | --- | --- | --- |
| Long | Long | Stable, 1 | Stable, 1 | High, $1 + \gamma m_d$ | High, $1 + \gamma m_d$ |
| Long | Short | Stable, 1 | Stable, 1 | High, $1 + \gamma m_d$ | High, $1 + \gamma m_d$ |
| Short | Long | Unstable, $1 + m_d$ | Stable, 1 | Low, 1 | High, $1 + \gamma m_d$ |
| Short | Short | Unstable, $1 + m_d$ | Unstable, $1 + m_d$ | Low, 1 | Low, 1 |

### The regulatory principle of protein expression dynamics under p53 pulsing

Let us examine the Hill type equation. For a very high binding affinity, *i*.*e. K*_*A*_ ≪*A*,

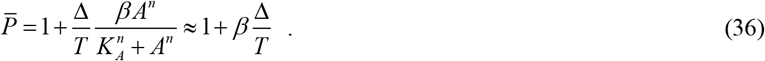

Therefore, the expression of target protein with high p53 DNA-binding affinity is insensitive to amplitude. Only through duration and frequency can fine-tuning of p21 protein expression be achieved, demonstrating the regulatory ability of duration and frequency beyond saturation (Fig 2). In addition, it is important to note that Δ *T* < 1, namely, the frequency must be less than that of 1 Δ. Thus, a situation in which too high a frequency causes protein expression to decrease is avoided. Furthermore, the results from the experiment showed that gene expression increased proportionally with TF frequency (35), which agrees with the prediction from the equation.

**Figure 2.**
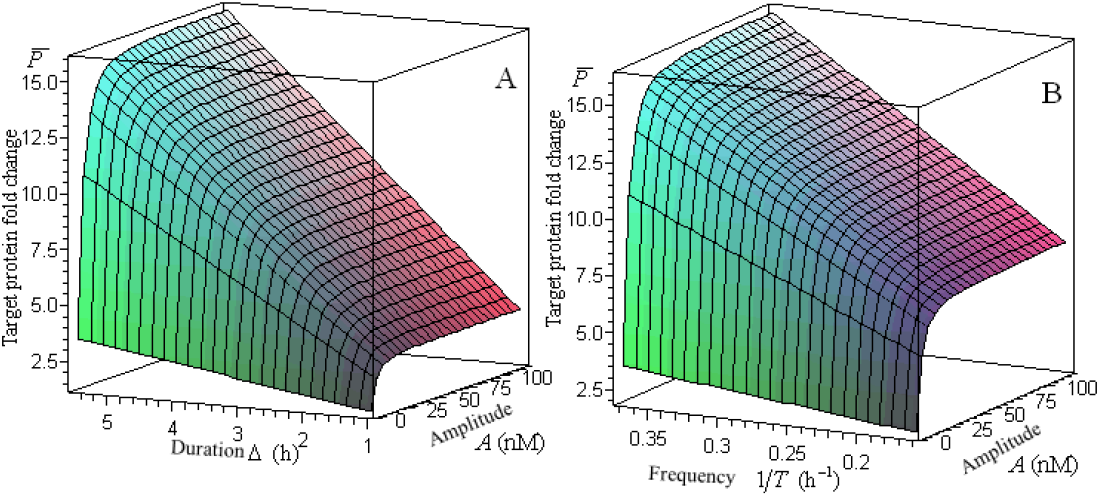
The effects of duration and frequency on p21 protein expression. Target protein expression nonlinearly reaches saturation with increased amplitude, however, it linearly increases beyond saturation as duration and frequency increase, thus, the cells achieved fine-tuning of gene expression using duration and frequency with high binding affinity. (A). *T* = 5.5 h, *K*_*A*_ = 4.9 nM, *n* = 1.8. (B).Δ = 2.75 h, *K*_*A*_ = 4.9 nM, *n* = 1.8.

The maximum mRNA expression of *β* was determined using binding affinity. The higher the affinity, the greater the maximal mRNA level (24, 34). For example, as shown in Table 1, for the *p21, GADD45, MDM2*, and *BAX* genes, *K*_*A*_ are 4.9 nM, 7.7 nM, 12.3 nM, and 73 nM (34), respectively, and the corresponding *β* values are 14.937, 9.157, 6.898, and 2.358, respectively (24). Therefore, the maximal gene expression level is mainly determined by the TF DNA-binding affinity, whereas the stability of gene expression is determined by the half-life of mRNA and protein. Therefore, the regulation of binding affinity has become a key issue in gene expression control. Gurdon et al. can greatly improve the binding affinity by extending the residence time of the TF target (36), allowing the duration and frequency of TF dynamics to finely tune gene expression.

### The fold changes in basal mRNA and protein expression are equivalent at steady state

For basal gene expression, let *S*(*t*) = 0, Equations 3 and 4, which describe the fold changes in mRNA and protein expression, respectively, become

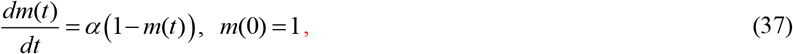

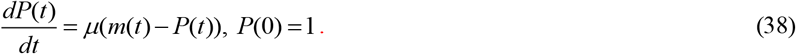

Therefore, at steady state the fold changes in basal mRNA and protein expression are

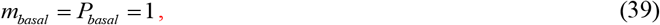

where *m*_*basal*_ and *P*_*basal*_ are the fold changes in basal mRNA and protein expression, respectively. By letting Δ= 0 or *A* = 0 in Hill-type equation 14, we can also obtain Equation 39.

Similarly, for basal gene expression, letting, Equations 1 and 2, which describe the concentrations of mRNA and protein expression, respectively, become

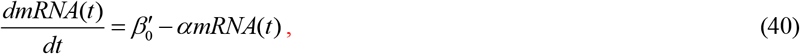

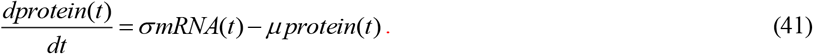

The steady state is

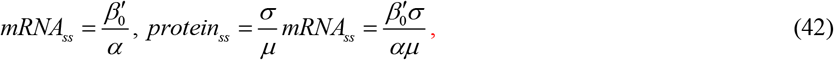

where *mRNA*_*ss*_ and *protein*_*ss*_ represent the concentrations of basal mRNA and protein at steady state, respectively. All the symbols and definitions used in this article are listed in Table 3. Therefore, for the basal gene expression system, the absolute levels of protein expression at steady state are determined by the rate constant of translation and degradation and mRNA levels. For any given gene, it must be noted that the above system has only one steady state.

**Table 3.**
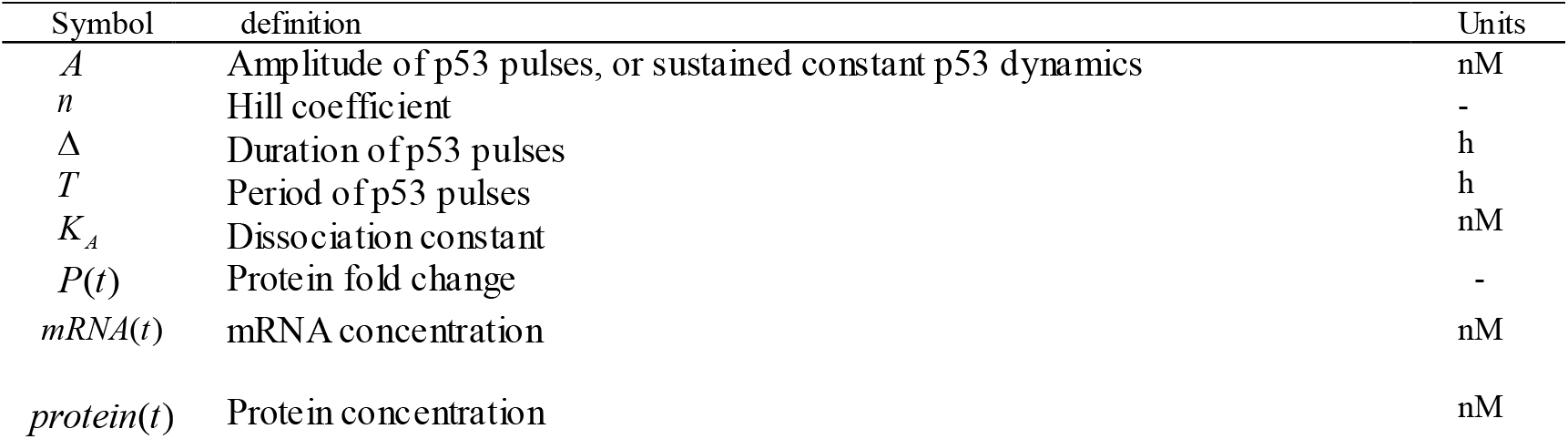
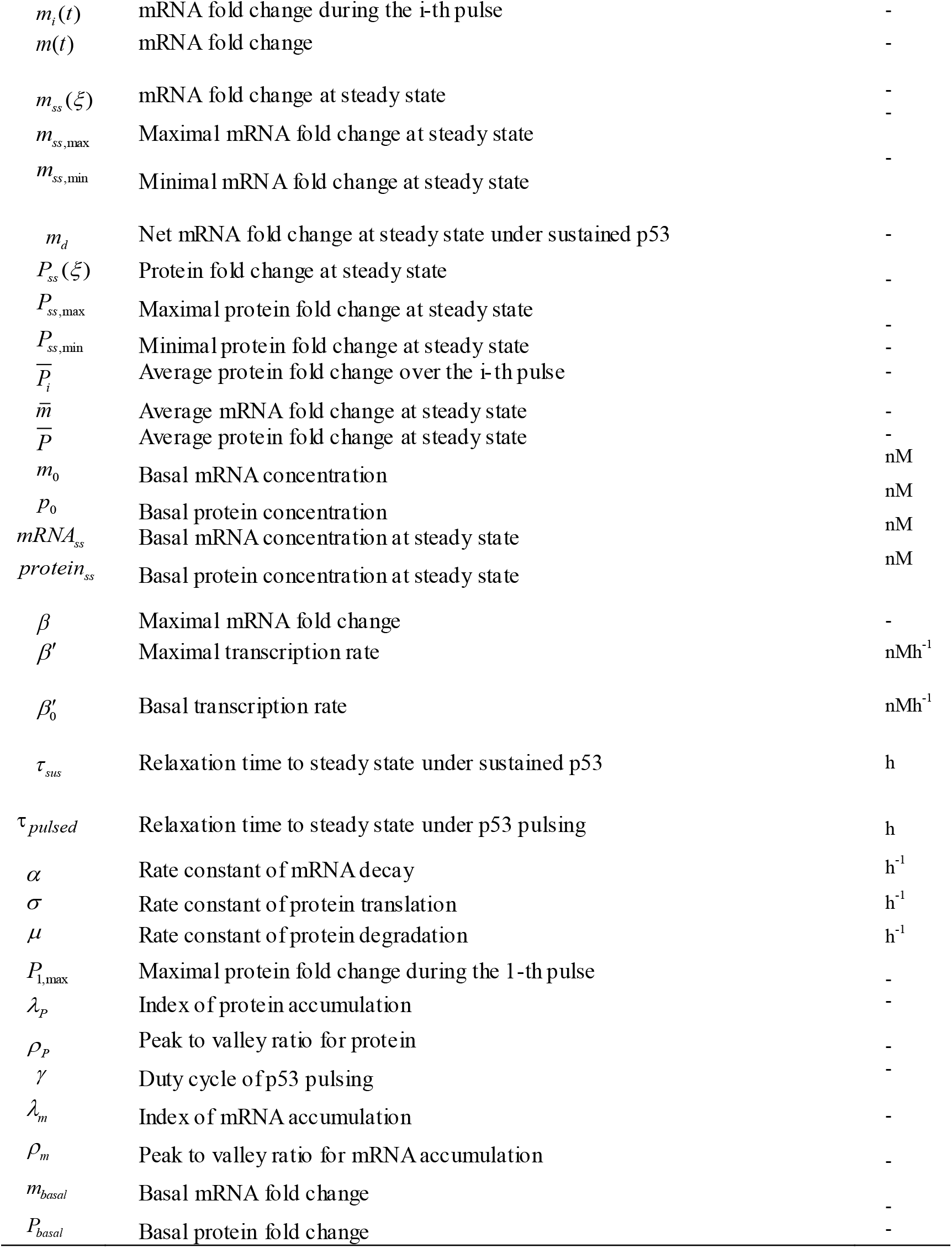
Variable and parameter definitions

| Symbol | definition | Units |
| --- | --- | --- |
| $A$ | Amplitude of p53 pulses, or sustained constant p53 dynamics | nM |
| $n$ | Hill coefficient | - |
| $\Delta$ | Duration of p53 pulses | h |
| $T$ | Period of p53 pulses | h |
| $K_A$ | Dissociation constant | nM |
| $P(t)$ | Protein fold change | - |
| $mRNA(t)$ | mRNA concentration | nM |
| $protein(t)$ | Protein concentration | nM |
| $m_i(t)$ | mRNA fold change during the $i$-th pulse | - |
| $m(t)$ | mRNA fold change | - |
| $m_{ss}(\xi)$ | mRNA fold change at steady state | - |
| $m_{ss,max}$ | Maximal mRNA fold change at steady state | - |
| $m_{ss,min}$ | Minimal mRNA fold change at steady state | - |
| $m_d$ | Net mRNA fold change at steady state under sustained p53 | - |
| $P_{ss}(\xi)$ | Protein fold change at steady state | - |
| $P_{ss,max}$ | Maximal protein fold change at steady state | - |
| $P_{ss,min}$ | Minimal protein fold change at steady state | - |
| $\overline{P}_i$ | Average protein fold change over the $i$-th pulse | - |
| $\overline{m}$ | Average mRNA fold change at steady state | - |
| $\overline{P}$ | Average protein fold change at steady state | - |
| $m_0$ | Basal mRNA concentration | nM |
| $P_0$ | Basal protein concentration | nM |
| $mRNA_{ss}$ | Basal mRNA concentration at steady state | nM |
| $protein_{ss}$ | Basal protein concentration at steady state | nM |
| $\beta$ | Maximal mRNA fold change | - |
| $\beta'$ | Maximal transcription rate | nMh<sup>-1</sup> |
| $\beta'_0$ | Basal transcription rate | nMh<sup>-1</sup> |
| $\tau_{sus}$ | Relaxation time to steady state under sustained p53 | h |
| $\tau_{pulsed}$ | Relaxation time to steady state under p53 pulsing | h |
| $\alpha$ | Rate constant of mRNA decay | h<sup>-1</sup> |
| $\sigma$ | Rate constant of protein translation | h<sup>-1</sup> |
| $\mu$ | Rate constant of protein degradation | h<sup>-1</sup> |
| $P_{1,max}$ | Maximal protein fold change during the 1-th pulse | - |
| $\lambda_p$ | Index of protein accumulation | - |
| $\rho_p$ | Peak to valley ratio for protein | - |
| $\gamma$ | Duty cycle of p53 pulsing | - |
| $\lambda_m$ | Index of mRNA accumulation | - |
| $\rho_m$ | Peak to valley ratio for mRNA accumulation | - |
| $m_{basal}$ | Basal mRNA fold change | - |
| $P_{basal}$ | Basal protein fold change | - |

The fold changes in basal mRNA and protein expression at steady state are

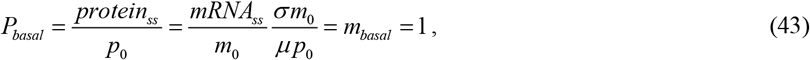

which is the same as Equation 39.

### Numerical results for MDM2 mRNA and protein expression

In order to verify the analytical results, as a example of p53 target gene *MDM2*, substituting available data into above analytical results, we can easily obtain the numerical results of MDM2 mRNA and protein expression. For gene *MDM2*, from Equation 9, the maximal fold change in MDM2 protein expression was

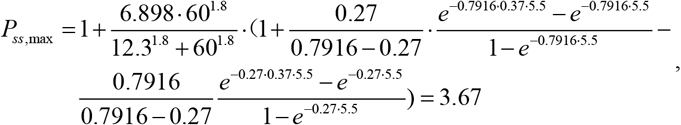

from Equation 10, the minimal fold change in MDM2 protein was

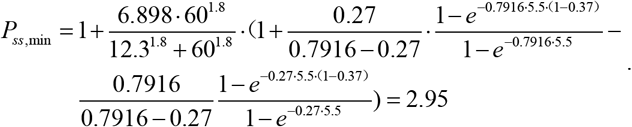

Thus, the peak-to-valley radio for MDM2 protein was

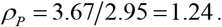

From Equation 26, the maximal fold change in MDM2 mRNA was

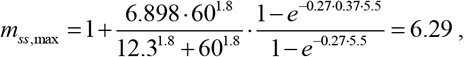

from Equation 27, the minimal fold change in MDM2 mRNA was

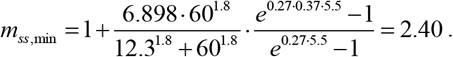

Similarly, the peak-to-valley ratio for MDM2 mRNA was

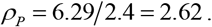

Considering that 1.24 < 2.62, therefore, although the fold changes in MDM2 mRNA and protein expression are both unstable, the degree of instability of MDM2 protein is much weaker.

Let us calculate the average MDM2 protein expression. From Equation 13, the average fold change over the 3-th pulse was

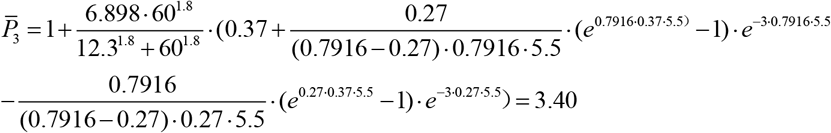

Similarly, the average fold changes over the 1-th and 2-th pulse were 2.63, 3.30, respectively. From Shi (26) Equation 19, the average fold change in MDM2 mRNA over the 3-th pulse was

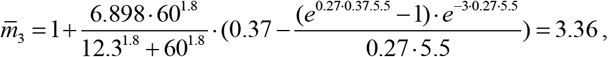

and 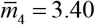. From Equation 14, the average fold changes in MDM2 mRNA and protein expression at steady state were

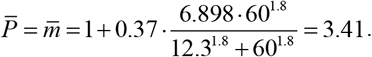

The observation values of MDM2 mRNA and protein were 2.2 or 2.0, 1.8, respectively. The deviation between prediction and observation is caused by two reasons. One is the instability of MDM2 mRNA and protein expression, the other is that the observation is over the cell populations rather than single cells.

## Discussion

TFs are nodes of a complex network, which is similar to a bridge connecting cellular signal transduction networks and transcription networks. Using the well-known TF p53 as an example, I proved that the steady-state fold changes in mRNA and target protein expression driven by p53 pulsing are the same. The Hill-type equation obtained may also be expected from previous methods (3, 11, 13, 16). Using this equation, we not only clearly understand the regulatory principle of gene expression, but also put this equation into practice. In addition, I provide two quantitative indicators to determine the degree of accumulation and stability of protein expression under multiple TF input pulses. These analytical results tell us that gene expression can be quantitatively predicted, what is more important, such analytical method is an effective way to solve problems, which may inspire us to solve more biological problems by this way.

For repressed gene expression, It was also proven that the fold changes in mRNA and target protein expression are the same at steady state, and a Hill-type equation with a negative coefficient can describe the fold change of target gene expression (37)(full text has not been submitted). Therefore, for both basal and activated or repressed expression, we may reach the consensus that the fold changes in mRNA and protein expression are the same at steady state and can be characterized by the Hill-type equation.

Both BAX mRNA and protein have very long half-lives, therefore, the fold changes in mRNA and protein become very stable, thus, the prediction from Hill-type equation agrees with observation well. The prediction in BAX mRNA and protein were both 1.35, the observation in BAX mRNA and protein were 1.40, 1.28, respectively, which come from the same lab under a radiomimetic drug. In addition, the observation value 1.72 in BAX mRNA is come from other lab under gama irradiation. The different stimulation may cause the deviation of observed values.

Hill-type equation reveals the regulatory principle of target protein expression in response to TFs pulsing. Amplitude modulation is effective with lower TFs-DNA binding affinity, however, for higher binding affinity, amplitude modulation quickly reaches saturation, duration and frequency can tune finely target protein expression beyond saturation. In addition, for a target gene, binding affinity, maximal mRNA fold change, mRNA and target protein half-lives can reflect the function of target protein, therefore, the complete observation data is necessary.

In Salazar et al. (32) and Shi et al. (25), for slow pulsing or fast dissociation, an equation similar to Equation 1 is introduced, and cannot be directly derived from the trajectory of the system. This may be an average steady state. In the limit of slow pulsing or fast dissociation, the peak-to-valley ratio tends to infinity. Thus, this average steady state is just zero(baseline). However, for fast pulsing or slow dissociation, another steady state can be derived not only in the trajectory but also in the average equation. Therefore, it is observable. In this case the peak-to-valley ratio is 1, so the system is stable. This equation is called the modified Hill equation which obtains from a ligand-receptor association and dissociation system under pulsing signals. Correspondingly, Equation 1 is called the Hill-type equation which comes from the synthesis and degradation of products upon oscillating signals (26). At present, there are only two types of Hill-type equations.

There are many models of gene expression. Some have been cited in previous research from TF to DNA to mRNA. Gene expression exhibits stochastic characteristics.(38-40). The impact of stochastic effects on gene expression has also been explained in the research from DNA to mRNA. The classic Hill equation has long been used in the modelling of biological systems (41-46). Mathematically, the Michaelis-Menten equation is equivalent to the Hill equation if the Hill coefficient is unity(47). The purpose of this study was to determine the principle of gene expression, and the results obtained needed to withstand practical testing, and the parameters of the results were measurable. Therefore, we can only flexibly develop a minimal model. Evidently, the basic purpose has been achieved.

Although gene expression dynamics are random and complex, the regulation principle is always deterministic and simple. “When a process depends on a range of different sources of randomness, instead of getting more complicated, it is possible for the different random factors to compensate for each other and produce more predictable results. Talagrand has given sharp quantitative estimates for this”(48). The Hill-type equation reflects the principle of deep simplicity of gene expression dynamics. The history of mechanics, physics, and chemistry indicates that the essence of nature is simple. Complex phenomena have evolved on the basis of the principle of simplicity. According to the regulation principle revealed by this equation, we can control gene expression from upstream of the genetic information flow described by the central dogma. Only four parameters need to be adjusted, namely, the duration, frequency, and amplitude of TF and binding affinity, and we can control the steady-state levels of mRNA and protein expression.

Because the levels of mRNA and protein expression at steady state are equal, and a large amount of transcriptome data has accumulated over the years; therefore, using this equation, we may predict the proteome simply by the transcriptome.

The classical Hill equation is applied when the TFs dynamics are constant. When Δ = *T* is applied, the equation is reduced to the classical Hill equation, therefore, this equation broadens the application of classic biochemical theory. Through the derivation process, the results represented by symbols are not only applicable to a specific TF, therefore, this generalized equation can be applied to any activated gene expression pathway driven by TF dynamics.

Hill wrote: ―My object was rather to see whether an equation of this type can satisfy all the observations, than to base any direct physical meaning on *n* and *K* (49, 50).‖ More than 100 years later, we have not forgotten his caveat (50).

## Appendix

### 1. The solution for Equation 5

The equation for target protein expression dynamics is

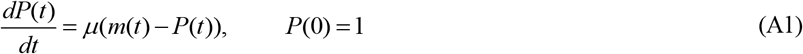

or

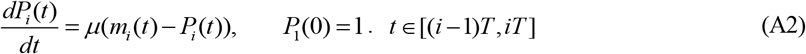

Denoting *ξ* = *t* − (*i* −1)*T* (32), then Equation A2 become

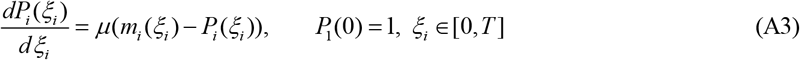

To solve Equation A3, letting general solution

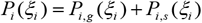

Then general solution to the homogeneous equation of Equation A3 is

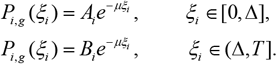

Assuming *α* ≠ *μ*, then a special solution for Equation A3 is

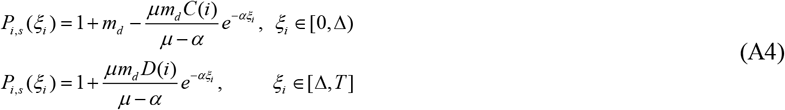

where

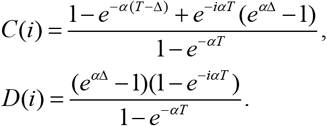

Thus, we can obtain the general solution for Equation A3:

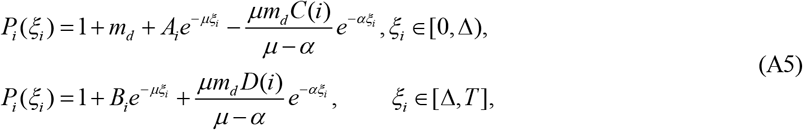

Using the condition (32)

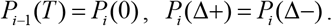

We can obtain the iterative equation

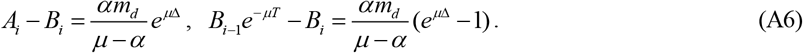

Considering *P*_1_ (0) = 1, thus,

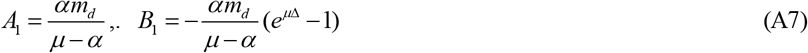

Denoting 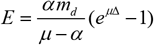, thus

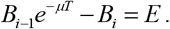

Thus,

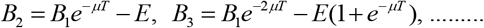

Generally, we have

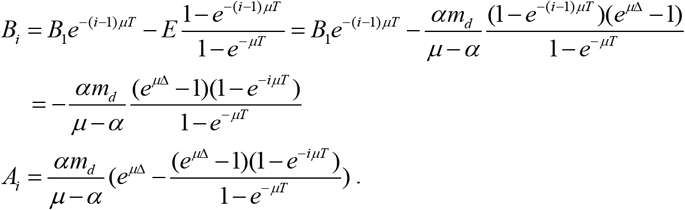

Thus, we have the solution Equation 7.

### 2. The average fold change of target protein expression

The average target protein expression levels is defined as(32)

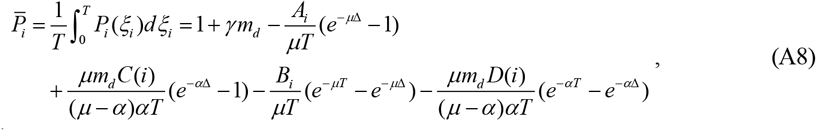

where

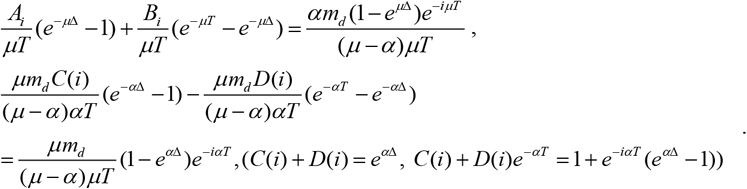

Thus

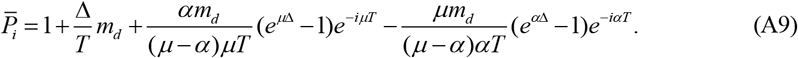

## Acknowledgements

I thank anonymous reviewers for several helpful comments.

## Ethics

This work did not require ethical approval from a human subject or animal committee.

## Data availability

The Data used in this article were obtained from [17, 23-26, 28, 33, 34] or are included in the text.

## Author contributions

X.S.: conceptualization, formal analysis, investigation, methodology, writing—original draft, writing—review, and editing.

